# A spatio-temporal model to reveal oscillator phenotypes in molecular clocks: Parameter estimation elucidates circadian gene transcription dynamics in single-cells

**DOI:** 10.1101/2021.08.04.455027

**Authors:** Måns Unosson, Marco Brancaccio, Michael Hastings, Adam M. Johansen, Bärbel Finkenstädt

## Abstract

We propose a stochastic distributed delay model together with a Markov random field prior and a measurement model for bioluminescence-reporting to analyse spatiotemporal gene expression in intact networks of cells. The model describes the oscillating time evolution of molecular mRNA counts through a negative transcriptional-translational feedback loop encoded in a chemical Langevin equation with a probabilistic delay distribution. The model is extended spatially by means of a multiplicative random effects model with a first order Markov random field prior distribution. Our methodology effectively separates intrinsic molecular noise, measurement noise, and extrinsic noise and phenotypic variation driving cell heterogeneity, while being amenable to parameter identification and inference. Based on the single-cell model we propose a novel computational stability analysis that allows us to infer two key characteristics, namely the robustness of the oscillations, i.e. whether the reaction network exhibits sustained or damped oscillations, and the profile of the regulation, i.e. whether the inhibition occurs over time in a more distributed versus a more direct manner, which affects the cells’ ability to phase-shift to new schedules. We show how insight into the spatio-temporal characteristics of the circadian feedback loop in the suprachiasmatic nucleus (SCN) can be gained by applying the methodology to bioluminescence-reported expression of the circadian core clock gene *Cry1* across mouse SCN tissue. We find that while (almost) all SCN neurons exhibit robust cell-autonomous oscillations, the parameters that are associated with the regulatory transcription profile give rise to a spatial division of the tissue between the central region whose oscillations are resilient to perturbation in the sense that they maintain a high degree of synchronicity, and the dorsal region which appears to phase shift in a more diversified way as a response to large perturbations and thus could be more amenable to entrainment.

## Introduction

Single-cell models typically fall into one of two categories: damped oscillators where rhythms arise through noise perturbation, so-called noise-induced oscillators, or limit cycle oscillators where the dynamics are subject to stable oscillations irrespective of the presence of noise [41]. In practice, and in the presence of multiple sources of noise, it is a difficult problem to differentiate empirically between noise-induced and robust limit cycle oscillators. As such, characterizing the spatial distribution of oscillatory phenotypes in dynamic gene expression processes remains an open problem. In the present paper we propose a novel spatially extended statistical methodology to perform parameter estimation and to assess the stability and empirical robustness of self-sustained oscillations at the single-cell level and hence to characterize oscillatory phenotypes of molecular clocks in a spatially distributed population of cells.

Biochemical reaction networks such as transcriptional and translational feedback loops (TTFLs) that give rise to oscillatory behaviour in gene expression and molecular clocks are complex, involving many chemical species and reactions. While the time evolution of molecular counts of chemical species in a reaction network can be described exactly by a Markov jump process (MJP), its complexity is in stark contrast to the availability of data as rarely can more than one species be observed at a time, and observations are typically obtained at discrete time-intervals as a result of experimental processes involving fluorescent reporter protein imaging. Model reductions of the full reaction network towards less parameter-intensive approaches that can feasibly be estimated from the experimental data are of considerable importance. The introduction of time-delays can approximate reaction events which are not of primary interest and thus tremendously reduce model complexity [28, 16, 23, 1]. Calderazzo et al. [5] propose the stochastic differential equation model with a distributed delay, where the natural variability in the delay time between reaction events is modelled by a probability distribution. The resulting stochastic representation provides a significantly reduced model which can realistically account for the intrinsic noise and rhythm generation inherent in the single-cell TTFL yet may feasibly be estimated from univariate experimental data. Despite model reduction, inference remains challenging due to the intractability of the transition densities of the MJP. Approximations in continuous state-space have been developed and are available when suitable assumptions on the system size hold, such as the chemical Langevin equation (CLE) [10, 16, 11] which aims at matching the infinitesimal mean and variance of the original MJP. Extensions to inference from observed data have been considered for the linear noise approximation (LNA)[22, 9, 34] which performs a linearisation leading to tractable Gaussian transition densities. The LNA has been extended to models with distributed delays [4], and a filtering algorithm which exploits the LNA was derived by [5] for likelihood-based inference, including Bayesian inference, for chemical reaction networks. In this study we extend the approach of [5] spatially through Bayesian hierarchical modelling of the parameters associated with the TTFL with a first order Markov random field prior distribution. This allows us to infer the parameters characterizing the TTFL across the tissue from existing experimental spatio-temporal gene expression data. We introduce the use of the robustness measure *V* to explore whether cells exhibit sustained or damped oscillations, and the inhibition profile (IP) to measure the relative impact of time-specific perturbations which also allows us to quantify entrainment characteristics of single-cell oscillators.

Recent interest has arisen in understanding the entrainment ability of the individual noisy neuronal circadian oscillators to form a precise biological clock [13] within the suprachiasmatic nucleus (SCN), the master clock of mammals, located in the hypothalamus of the brain. The circadian rhythms generated by the TTFL in SCN neurons are essential to coordinate daily patterns of physiology and enable entrainment of behavioural rhythms to environmental cues [17]. Accurate timekeeping is an emergent property of the SCN, relying on the synchronization of the circadian oscillations of approximately 20,000 neurons. At the single-cell level, these constitute stochastic clocks which synchronize through a combination of firing and rhythmic neurotransmitter release to form a biological clock which is precise overall [40]. Intrinsic noise arising from the low copy numbers of molecules [8] plays a significant role in the heterogeneity of neuronal rhythms and possibly in the generation of stable circadian rhythms [41]. An application to bioluminescence-reported expression of the core clock gene *Cry1* across organotypic mouse SCN provides an ideal test-bed of our methodology to study the emergent spatio-temporal circadian oscillatory dynamics of gene expression in SCN neurons.

## Results

We propose a mathematical model that can be fitted to spatio-temporal expression data of a single gene. To ensure reproducibility we studied three experimental replicates, labelled Rep.1–3, of CRY1-LUC measured across organotypic mouse SCN tissue over 5–6 days.

### A spatially extended distributed delay model with measurement equation

The model integrates three stochastic components (i) the intrinsic molecular noise of the dynamic TTFL processes at single-cell level, (ii) a measurement model to incorporate both measurement noise and signal aggregation over time of camera exposure, and (iii) a spatial model that captures heterogeneity in the spatial distribution of the transcriptional dynamics.

#### (i). The TTFL with distributed delay

To describe the stochastic dynamic TTFL transcription processes at the single-cell level we assume that mRNA abundance at time *t* and location *i, X*_*i*_(*t*), is the result of a stochastic birth and death processes that can be approximated by a CLE of the form [16, 5]

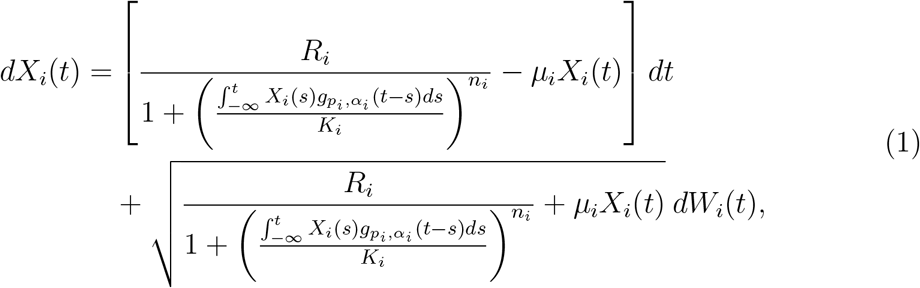

where *W*_*i*_(*t*) are standard Wiener processes. Gene transcription is classically modelled by a Hill-type transcription function [12] with maximum transcription rate *R*_*i*_, Hill coefficient *n*_*i*_, dissociation coefficient *K*_*i*_, while mRNA degradation is assumed to occur linearly with rate *µ*_*i*_. The term 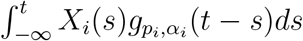 i.e. the expected mRNA available at *t* over a distribution of delay times, serves as a transcription factor proxy where we approximate the delay distribution, 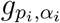, by a Gamma density with shape and rate parameters *p*_*i*_ and *a*_*i*_, respectively, which in practice is truncated at *τ*_max_ for computational tractability. For our data we set *τ*_max_ = 24h.

#### (ii). Measurement model for bioluminescence reported gene expression

The following form for the measurement model is assumed for light intensity of bioluminescence-reported gene expression data at location *i* at time point *t* ′ *∈* ℕ, *Y*_*i,t′*_[5]

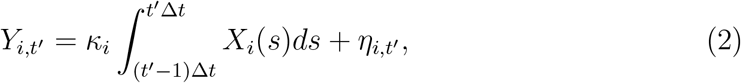

where *κ*_*i*_ is a scaling factor between the recorded light signal emitted by a reporter protein and mRNA, which may be assumed to vary spatially, for instance due to an irregular geometry of the biological sample. The limits of integration are determined by the exposure time interval setting of the camera, e.g. Δ*t* = 0.5h for our data, and *η*_*i,t′*_ is a Gaussian zero-mean random variable that represents measurement noise which may have a location specific variance.

#### (iii). Spatial model

We assume a simple but general multiplicative random effects model where a parameter value at location *i* is given by the overall spatial mean, *θ*, perturbed by a location-specific random effect described by exp *ϵ*_*i*_ such that

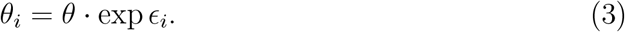

This model is assumed for each of the parameters *R*_*i*_, *K*_*i*_, *n*_*i*_, as well as the mean and standard deviation of the delay distribution. The assumed neighbourhood structure implies a first-order Markov random field, specifically that the random effect for a given parameter at location *i* is a priori dependent on the random effects of the same parameter associated with the 8 surrounding locations. This spatial model captures dependence between neighbouring locations only through a similarity between the parameters of the TTFL without making any mechanistic assumptions about the nature of cell communication where joint timekeeping is partially emergent due to network properties. While the advantage is that no further non-identifiable parameters were introduced we note that our parameter estimates of the TTFL will adjust necessarily to integrate these processes.

### Parameter estimation and model fit

As the posterior distribution resulting from components (i)-(iii) is intractable, we design a Markov chain Monte Carlo (MCMC) algorithm to draw a large number of samples from it (see Materials and methods). The posterior means of the five parameters associated with the Hill-function are shown in Fig.1. We find a pronounced spatial structure with regards to Hill-coefficient, *n*, and dispersion of the delay distribution, for all three replicates. Central locations of the tissue samples exhibit higher estimates of both parameters while edges exhibit lower estimates. These two parameters are important in determining the dynamics of the oscillator model in Eq. (1) [38], which is investigated further in Fig.4 and 6. We note that the delay dispersion can be equivalently summarized by the entropy of the delay distribution

**Figure 1:**
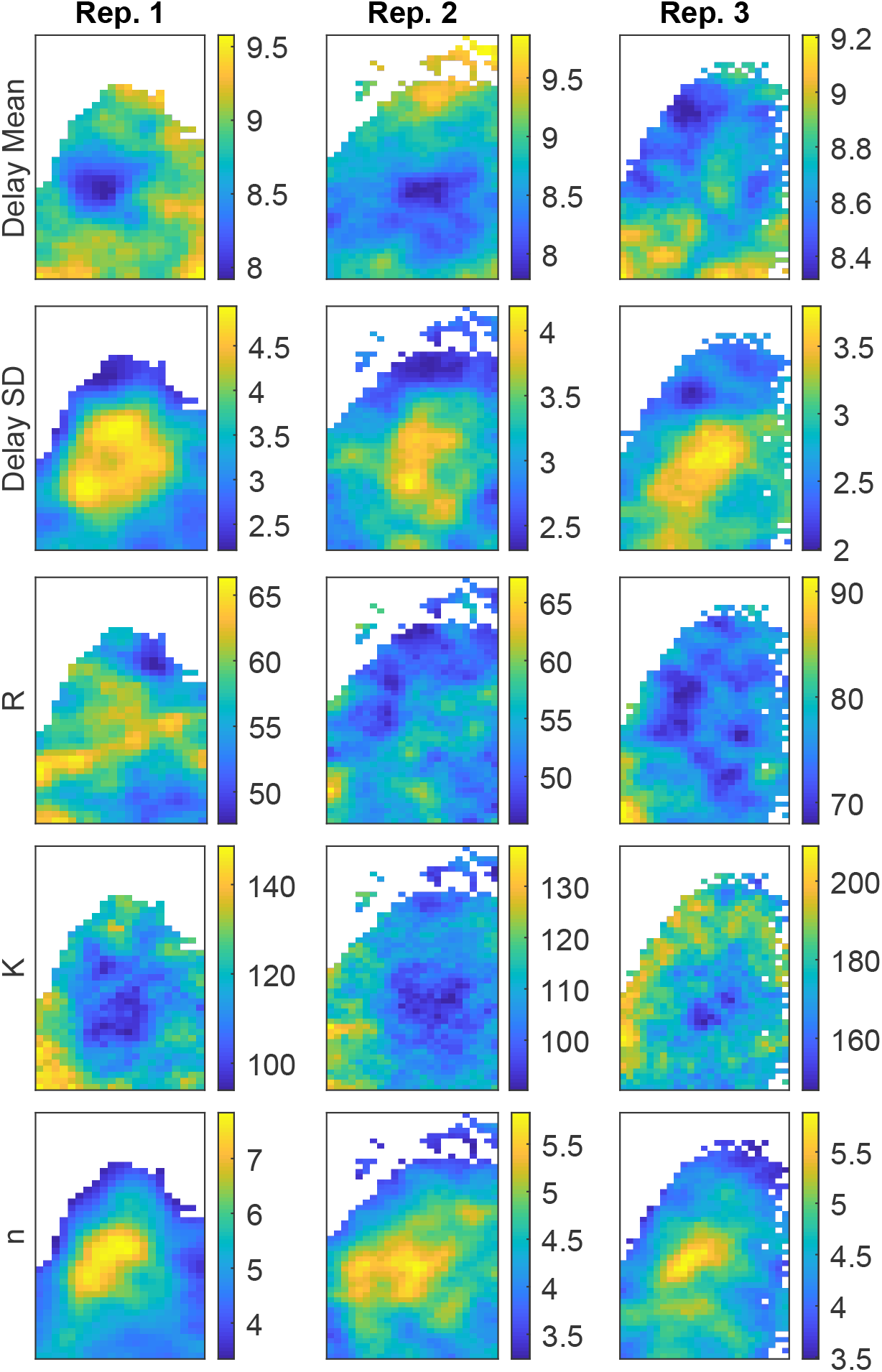
Spatial distribution of posterior means for delay mean (*µ*_Γ_), delay SD (*σ*_Γ_), maximum transcription rate (*R*), dissociation coefficient (*K*) and Hill coefficient (*n*).

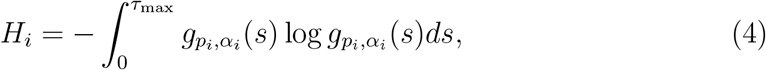

which provides a measure of the information contained in the delay distribution.

The maximum transcription rate *R* (molecules/h) and dissociation coefficient *K*, which both relate to the scale of molecular population sizes, exhibit low systematic spatial variation across the three replicates. The posterior means are overall similar in scale for replicates 1 and 2, while somewhat higher in replicate 3, suggesting that the degree of molecular noise, which informs the population size estimates via the Gaussian approximation, is homogeneous across the organ and consistent across replicates. The estimated mean delay times for the *Cry1* molecular oscillators are around 8-10 hours with a delay dispersion (SD) of 2-5 hours. The mean of the delay distribution tunes the period of oscillations produced by the model in Eq (1), and while exploratory spectral analysis of data suggest close to 24h periodicity across all tissue for all three replicates, the estimates of the model parameter exhibit some spatial variability. A presumptive cause of variability in the estimates is that the period of oscillations is not fully captured by the delay mean due to the fact that the asymmetry of the gamma distribution varies with the dispersion parameter (see Fig.6), hence the implied period of oscillations may have a lower variability than the mean estimates in Fig.1 as opposed to in a fixed delay model, which is a limiting case of Eq (1) as 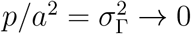.

To evaluate model fit, location-specific residual time series are obtained by computing the differences between the observed data and predictions produced by the estimated local model filter output. The outcome of the Kolmogorov-Smirnov test (with Bonferroni correction) at each of the 840, 1152, and 1125 locations of the three replicates accepted the null of Gaussianity of the residuals for all except a mere 5 locations (i.e. 0.16% of those considered). Residual time series were evaluated for further periodicity by means of a spectral bootstrap analysis [6]. Fig.2 shows the mean period at locations which have at least one period in the range 1-30h. We note that since all residual series are free of 24-hour periodicity the fitted model successfully explains the stochastic circadian dynamics observed at single-cell level. It appears that SCN tissue locations tend to exhibit an additional, albeit small, 12-hour residual periodicity. Evidence of gene expression with 12-hour periodicity has previously been found in hundreds of genes, e.g. in mouse liver tissue [18, 43].

**Figure 2:**
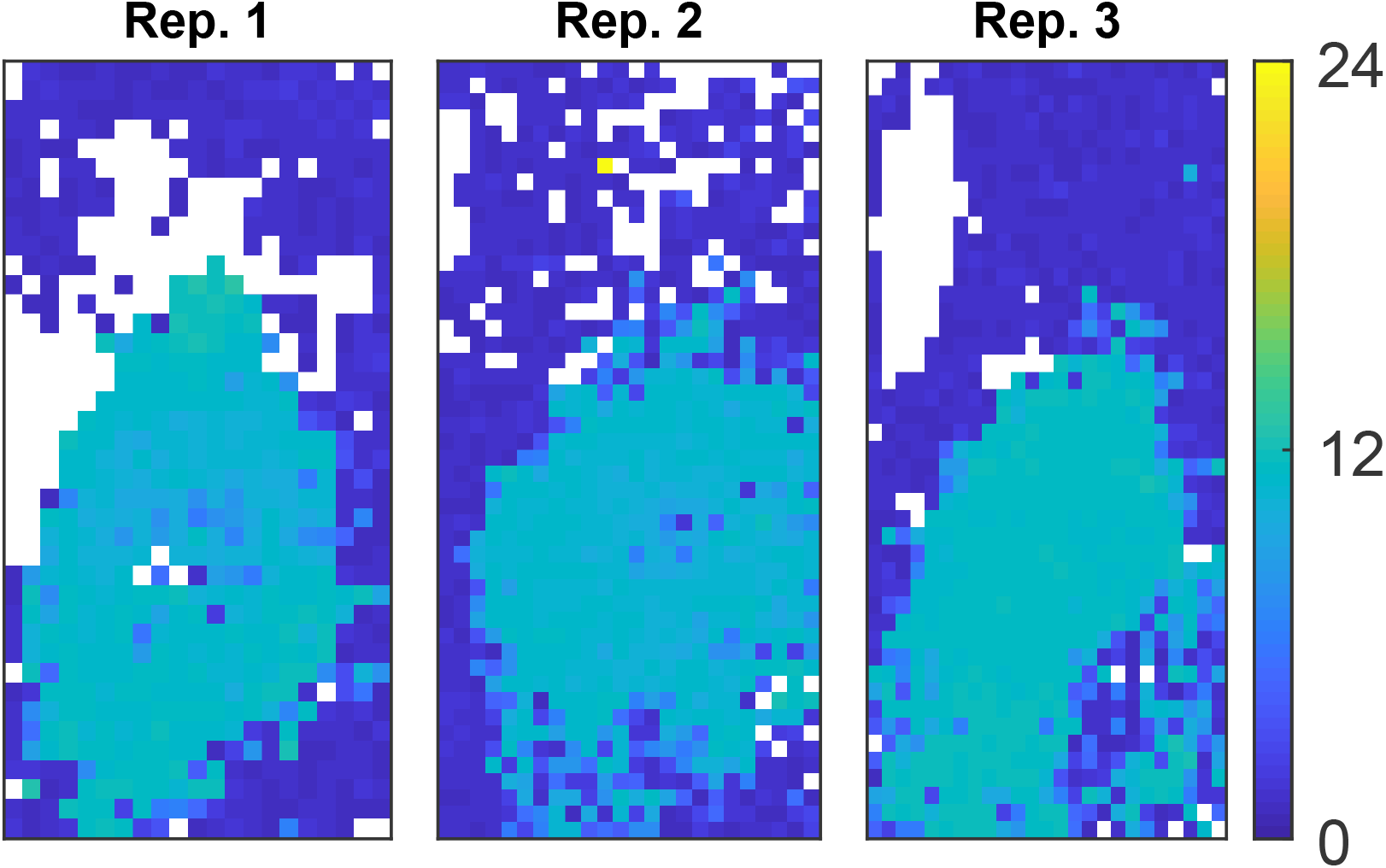
Spatial distribution of residual periodicity for locations where 99 % spectral bootstrap confidence intervals have an endpoint in the range 1-30h. Residuals are mostly free of 24 hour periodicity while locations corresponding to SCN tissue typically exhibit additional low-amplitude 12-hour periodicity.

### Estimating robustness of oscillations and regulatory responsiveness

Using a general definition of robustness *V* of system *s* with regard to property *a* evaluated by function 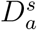 against a set of perturbations *P* with distribution *π*_*P*_ by [20]

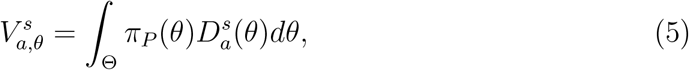

we propose an estimator of robustness where *π*_*P*_ is taken to be the Bayesian posterior distribution of parameters. With this choice of *π*_*P*_, empirical robustness captures parameter uncertainty along with its implications on some objective characteristic of interest of the system. This coincides with the Bayesian posterior probability of property *a* as measured by function *D*. Hence the evaluation function *D*(*θ*) takes the value 1 if *θ* implies limit cycle oscillations of the deterministic mean of the model in Eq (1) and 0 if the oscillations are damped (see Materials and methods, Macroscopic stability for further details on the stability criteria). Noting that Eq (5) is an expectation of 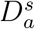 a posterior mean estimate of 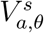 can be computed by evaluating 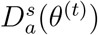 at each iteration *t* of the MCMC algorithm and averaging the resulting chain.

Furthermore, to quantify the responsiveness of transcriptional regulation to the inhibiting species, we introduce the Inhibition Profile (IP) as the (negative) gradient of the transcription rate with respect to the past levels of mRNA. The result is evaluated at and divided by the equilibrium solution, *x*^*∗*^, to obtain the effect at an average population size and a function that does not scale with population size. The IP is thus closely related to the phase-response curve, typically used to study entrainment of circadian oscillators to external stimuli [19], where instead of relating extraneous perturbations at different times of the cycle to phase shifts, here the IP gives the delayed marginal change in transcription rate induced by perturbations across the cycle. Due to the intrinsic noise-term of the model in Eq (1), the associated phase shifts are stochastic and we explore their distributions further in a simulation study (see Fig.7). The marginal inhibition due to a perturbation at *k* [0 ∈ *τ*_max_)time ago is given by

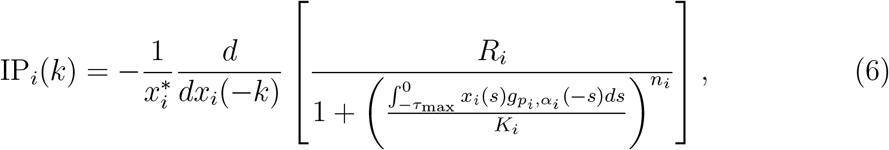

and the resulting function of *k* measures the relative effect of a small perturbation of mRNA in the preceding cycle on the transcription rate at time *t* = 0.

### Results for *Cry1* oscillatory phenotypes in mice SCN

The estimated oscillator robustness allows us to identify distinct regions (see Fig.3) of *Cry1* oscillatory phenotypes, firstly between the SCN tissue, which exclusively exhibits a high degree of robustness associated with strong limit cycle dynamics, and the immediately surrounding tissue exhibiting low empirical robustness or ‘noisy oscillators’. This result is very important as it confirms that the proposed modelling approach manages to correctly distinguish the robustness of the clock within the SCN from the non-robust or noisy oscillatory behaviour outside or at the edge of the tissue, noting that data from both regions appear indistinguishable to the eye or even standard exploratory methods such as spectral analysis. Within the SCN a small number of locations in some central and ventral clusters of Replicates 1 and 2 appear to exhibit weak oscillations or substantial uncertainty. This finding may be related to spatial variation of neuropeptide expression profiles as additional analyses, not reported here, of *Cry1* expression from VIP-depleted SCN [27] using our methodology produces robustness estimates that are consistent with noise-induced oscillations across all imaged tissue, i.e. 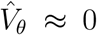 A multivariate analysis of all estimated TTFL parameters indicates that robustness of oscillations within the SCN is predominantly determined by three parameters, namely the Hill-coefficient, *n*, the dispersion or entropy of the delay distribution, 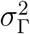, and the degradation rate, *µ*. Fig.4 shows that locations with high empirical robustness are found along a line of positive trade-off between the two parameters that are associated with the transcriptional regulation within the TTFL, i.e. the Hill-coefficient, with estimated values between 3 and 8, and delay dispersion whose estimated values are between 2 and 5 hours. With all other parameters held constant, we can see that an increase in the delay dispersion or entropy *reduces* the robustness or propensity for limit cycle dynamics. Under the assumed model this implies that a higher Hill-coefficient can compensate for a higher delay dispersion to obtain a robust oscillator phenotype.

**Figure 3:**
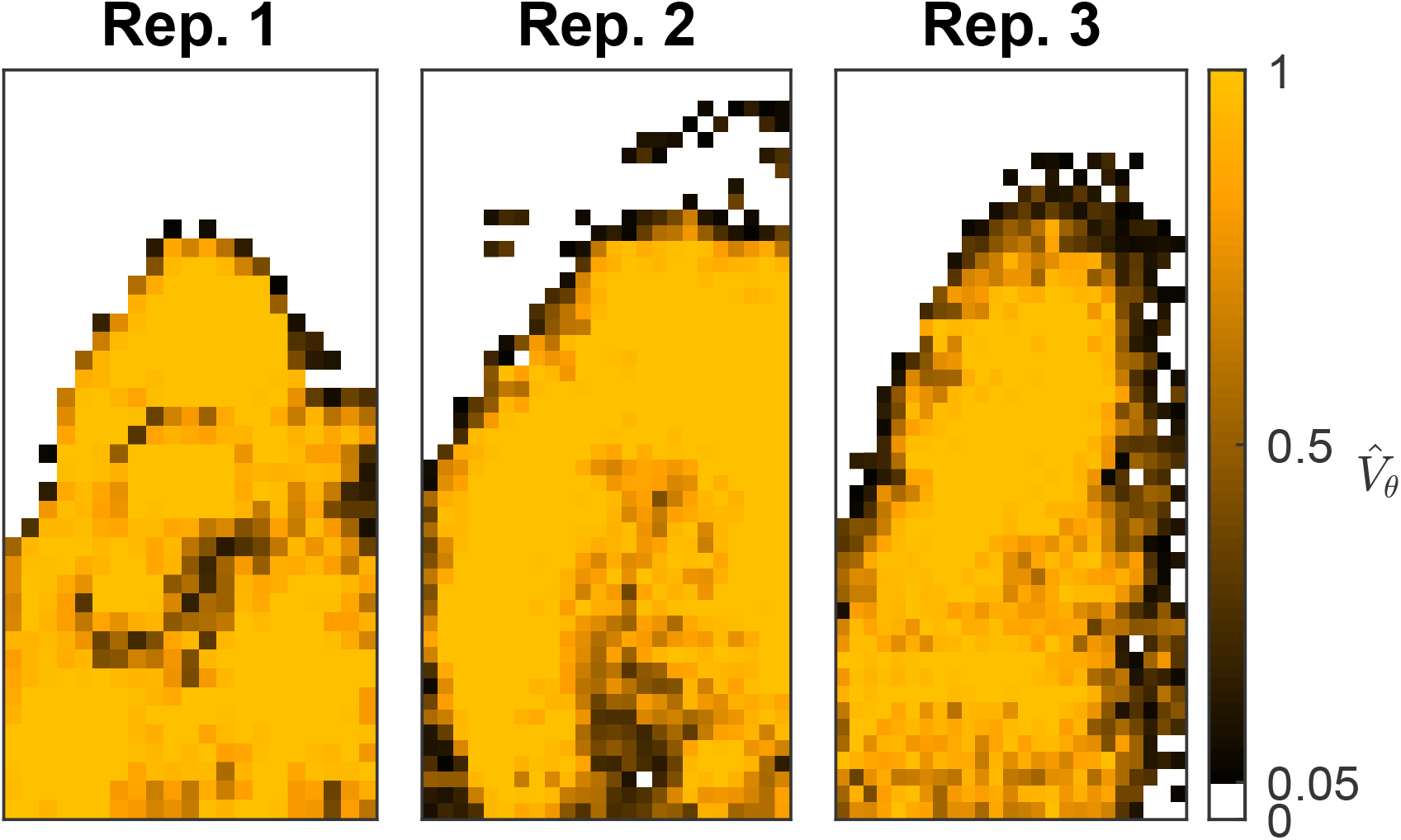
Spatial distribution of oscillator robustness, 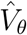 for three experimental replicates of CRY1-LUC data. A cut-off of 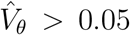 correctly differentiates the SCN from surrounding tissues. Most SCN tissues exhibit robust oscillations through limit cycle dynamics of the deterministic mean of the model 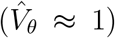 while the surrounding tissues exhibit noise induced oscillations 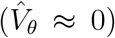. A few locations, found along the SCN edges and in some central and ventral clusters of Replicate 1 and 2, exhibit weak oscillations or substantial uncertainty 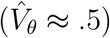

**Figure 4:**
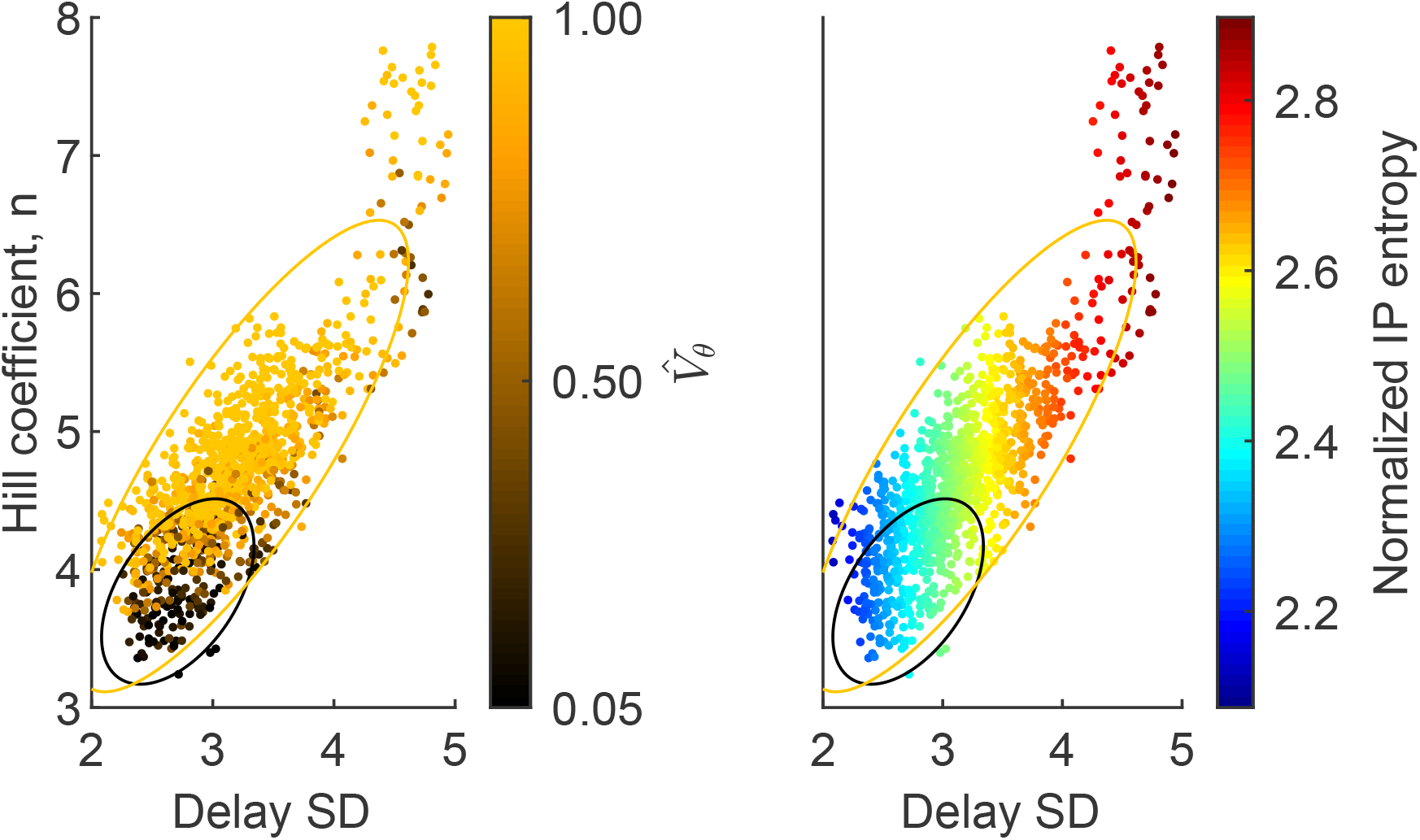
Scatter plot of posterior means of Hill coefficient and delay distribution dispersion (standard deviation) for three experimental replicates. Locations are sub sampled for visibility. In the left panel color is scaled by the estimated robustness of oscillations. Locations with high empirical robustness are found along a ridge formed by the Hill-coefficient and delay dispersion while locations closer to the cutoff 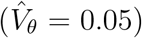 exhibit lower estimates of both parameters. In the right panel color is scaled by the entropy of the standardized IP. Ellipses represent the 95 percent contours of bivariate Gaussian distributions with means and covariance matrices of sub-populations defined by 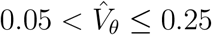 (black) and 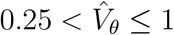 (yellow).

### Differential responsiveness of circadian oscillators in SCN

It appears that the IPs are more concentrated around a larger maximum for oscillators with a lower delay dispersion suggesting that, given some mean delay time, the inhibition happens more directly over a shorter time span than for an oscillator associated with a wider IP. This is tuned by the entropy of the delay distribution where oscillators with a low delay entropy, are subject to inhibitory dynamics that resemble a step function, or a fast transcriptional ‘on-off’ switch, whereas the inhibitory effect is spread over a longer time scale in oscillators associated with a larger entropy.

Furthermore, the total inhibition, defined as the integral of the inhibition profile IP with respect to *k*, is typically greater for the wider IP profiles due to the higher Hill coefficient. To visualize their spatial distribution, the IPs are summarized by their location-specific entropy for the three experimental replicates in Fig.5. The data show that oscillators with higher entropy are located in the central SCN, while low-entropy IPs are predominantly found in the dorsal tissues. As it has been found that the ventral SCN has a higher density of intercellular connectivity compared to the dorsal [36], and re-entrainment characteristics of SCN neurons following light-induced jet lag suggests higher proportions of fast and slow-shifting cells in the dorsal and ventral SCN, respectively [7], we hypothesize that the spatial localization of the information content of the IPs is indicative of spatially different entrainment characteristics.

**Figure 5:**
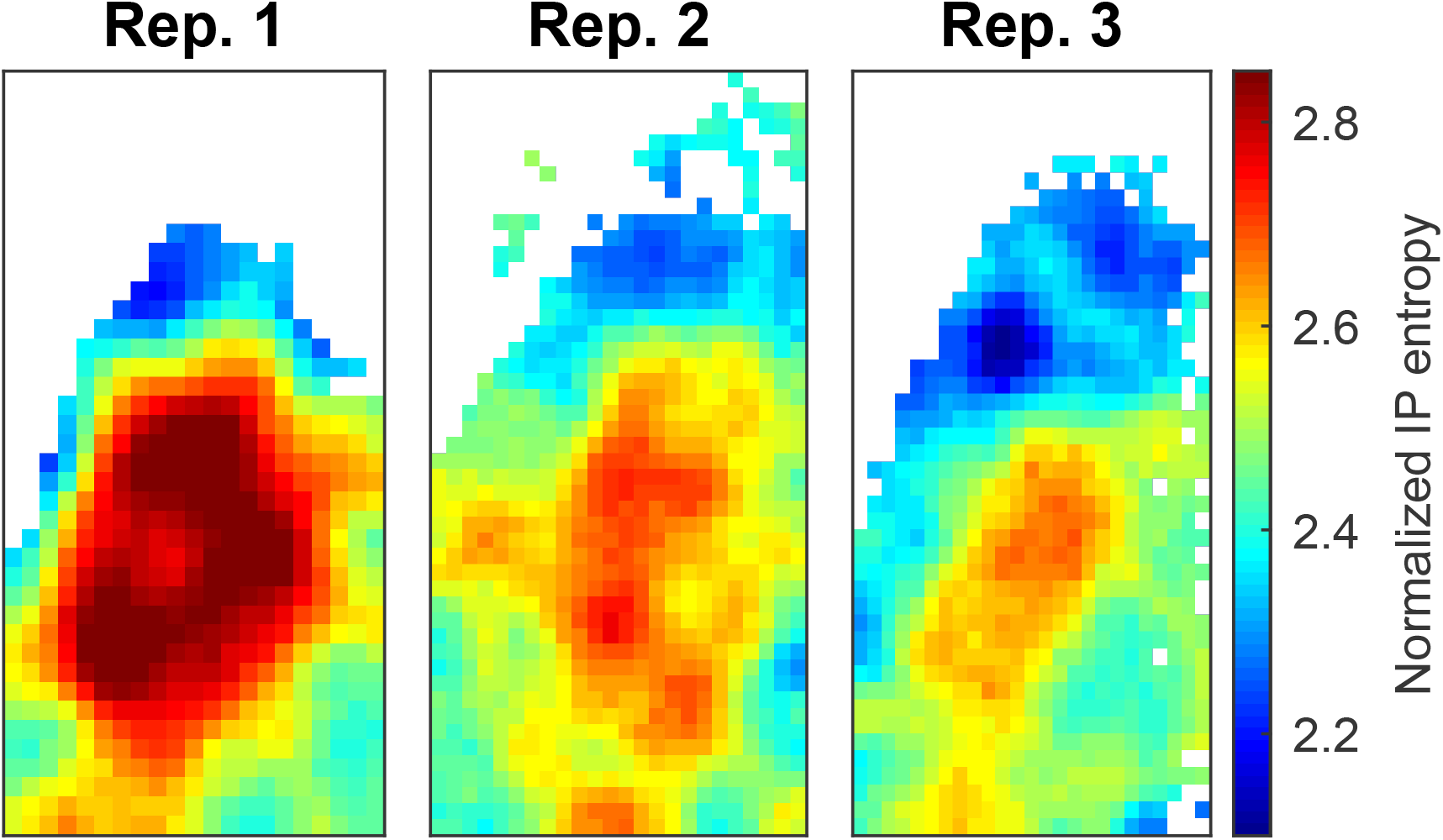
Entropy of the standardized IPs across the three experimental replicates. Wide IPs (high entropy) are found predominantly in central SCN while narrow IPs (low entropy) are localized to the dorsal SCN.

### Differential entrainability of circadian oscillators in SCN

To investigate this hypothesis we conduct a simulation study to compare the phase distributions of two prototypical oscillator types associated with two points located on the trade-off ‘ridge’. Both types exhibit robust limit cycle dynamics but are at opposite ends of the spectrum regarding their entropy. For simplicity, we shall refer to the oscillator corresponding to the lower point, characterized by a smaller *n* and delay entropy, as ‘Type I’, and to the other point as ‘Type II’, noting that Type I oscillators have IPs corresponding to dorsal SCN while Type II are representative of central SCN neurons. While no explicit synchronizing mechanism is assumed in our model, we define synchrony as the degree with which trajectories of the stochastic oscillators stay synchronized after a common initial condition. Analyzing entrainment and synchronization by studying ensembles of oscillators is discussed in [37]. We place an implicit assumption on the network topology that each individual oscillator has identical exposure to a given perturbation, while the response is driven by the cell-autonomous stochastic mechanisms (see [14]). Ensembles of gene expression trajectories from the two types of oscillators are forward simulated from the stochastic delayed differential equation model in (1) using an Euler-Maruyama approximation and we examine the effect of perturbations on the phase distribution for each ensemble. The perturbations are specified as positive shocks in the concentration of mRNA with an amplitude consistent with a typical estimated peak concentration (150–200 molecules) and varying duration (1–4 hours) thus controlling the overall size of the shock. While such sudden increases in mRNA are inconsistent with intrinsic noise in the single negative feedback loop in (1) where transcriptional regulation by Reverb/RORA loops is assumed constant and captured by *R*_*i*_, there is evidence that exogenous shocks, e.g. light, may dissociate interacting intracellular feedback loops [31]. Hence, the perturbations are considered deviations of varying magnitude from the single-cell’s unperturbed oscillation. The effect of perturbations of four increasing sizes on the ensemble’s synchrony is shown in Fig.7 along with polar histograms of the resulting phase distribution relative to the unperturbed trajectories, estimated from the subsequent five cycles.

**Figure 6:**
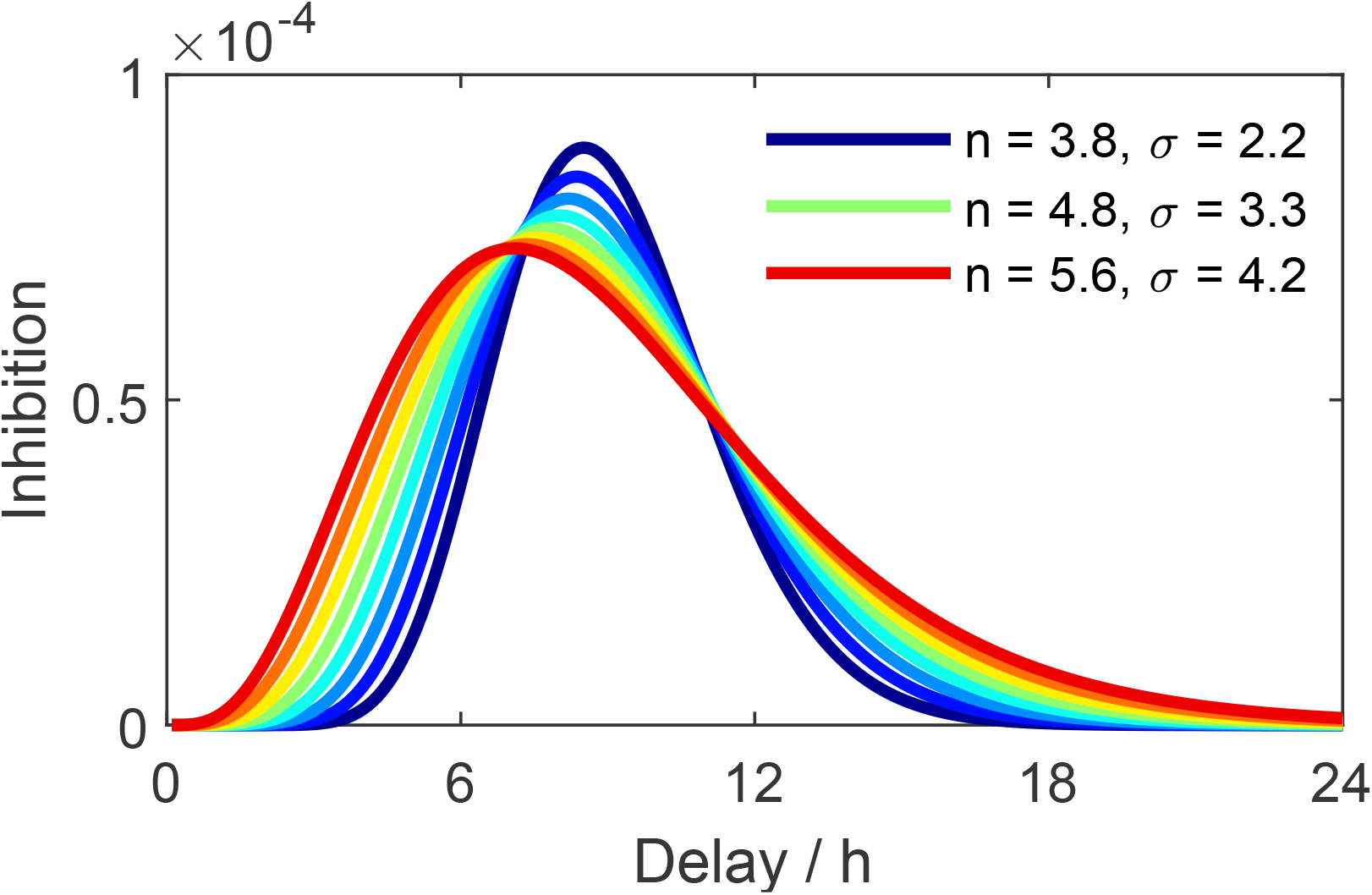
Inhibition profiles, calculated as the negative gradient of the transcription rate with respect to the initial data divided by the equilibrium population size, for parameter combinations along the ridge formed by the Hill-coefficient and delay dispersion. A high delay dispersion (red) distributes the inhibitory response over a longer time scale, while for low delay dispersion (blue) the inhibitory response is concentrated around the mean of the delay distribution.

**Figure 7:**
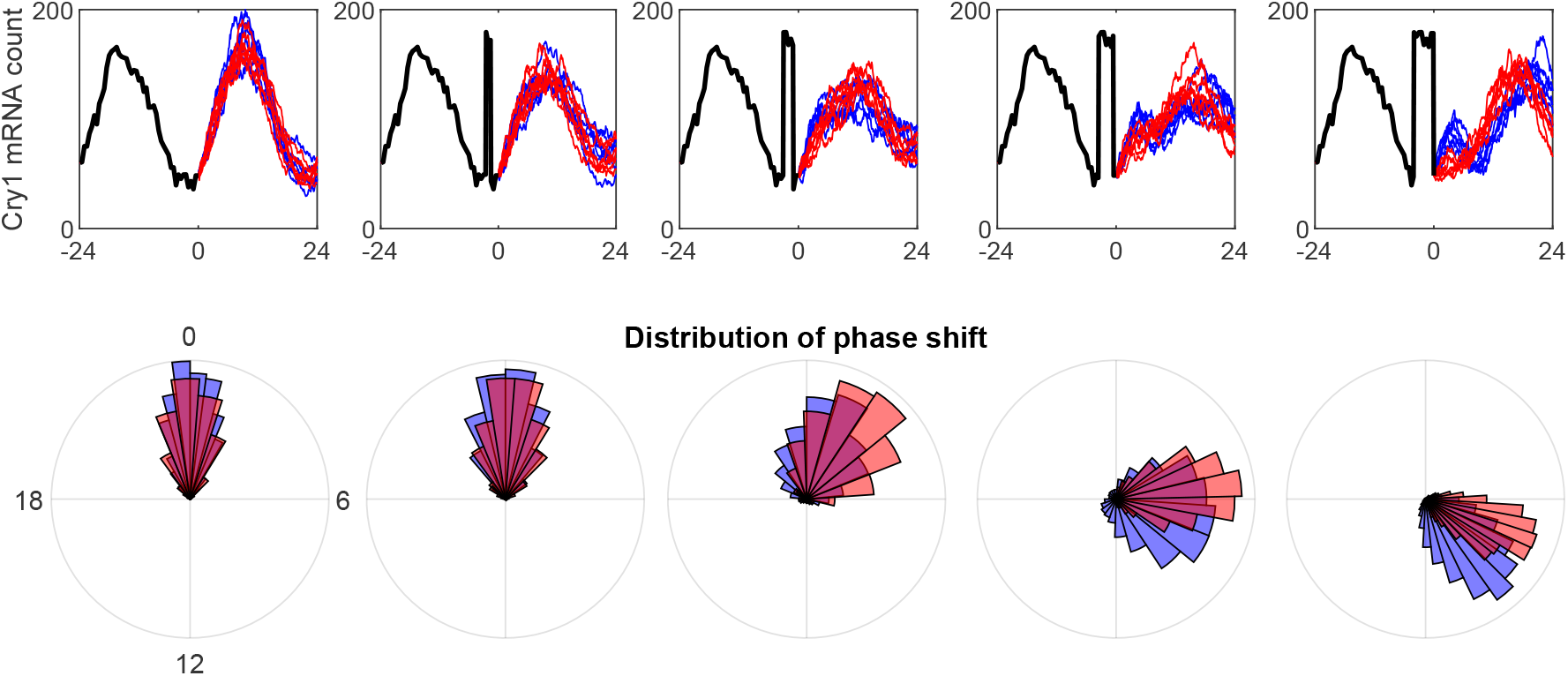
Differential responses to large positive perturbations of varying duration. Parameters are set to *R*_0_ = 50, *K* = 100, *µ* = 0.25 for both oscillator types while the Hill coefficient is set to *n* = 5.57 and *n* = 3.82 respectively. Delay mean is set to 9.4 and 9.1, and delay standard deviation is set to 4.2 and 2.2 respectively. Initial data is obtained by scaling the light signal by *κ* = 2.5×10^−3^ which is a typical estimate for SCN tissue. Perturbations are defined as a positive shock of 130 molecules during the timing of the minimum Cry1 concentration in the initial data. Trajectories are simulated using an Euler-Maruyama approximation to the model in (1) with time step set to *dt* = 0.1h. Bootstrap phase distributions are obtained by simulating 2000 paths and calculating the phase of the subsequent five cycles after the visualized paths in the figure.

For unperturbed trajectories it is verified that we reconstruct the reference case, namely that both oscillator types maintain synchronized ensembles during the whole simulation period of 6 days. The smallest perturbation duration (1 hour) induces a similar increase in the phase dispersion for both types but does not produce a phase shift in either oscillator type. As the perturbation duration increases to 3 hours the synchronization ability of both types starts to differentiate in that it deteriorates for Type I oscillators, while Type II oscillators remain relatively synchronised and perform a smaller phase-shift. In the largest perturbation case, Type I oscillators regain synchrony to a new circadian regime and are subject to a phase shift that is significantly larger than that of the Type II oscillators. These findings suggest that ensembles of oscillators of Type II, found in the central SCN neurons, remain relatively synchronised when subjected to large perturbations of varying duration, and phase-shift jointly to a new circadian regime in a predictable fashion. On the other hand, Type I dorsal oscillators may exhibit the same degree of synchronisation when unperturbed, but we have demonstrated that the ensemble’s time-keeping was disrupted by a sufficiently large perturbation.

The difference in response of the two oscillator types to the largest perturbation highlights a trade-off between synchrony and entrainability: while ensembles from Type II central SCN oscillators phase-shift in unison, their shift is smaller compared to the dorsal Type I oscillators. The total inhibition, i.e. the magnitude of the integral of the inhibition profile, is typically greater for the central neurons due to the higher value of the Hill coefficient, suggesting that they experience a higher degree of inhibitory pressure. While not modelled explicitly here, this may be indicative of inter-cellular processes where inhibition mechanisms are in part shared between neighbouring cells. Slight phase differences between connected neurons combined with shared inhibition may lengthen the time scale at which small increases in TTFL species contribute to transcriptional repression, consistent with our findings. Dorsal cells are less densely connected and as such, single-cell mechanisms that act under a shorter time scale may play a greater role in their rhythm generation.

From a modelling point of view, ensemble synchronization of centrally located oscillators can be understood by the delay distribution acting as a regulator of delayed Cry1 mRNA. A higher delay dispersion represents a larger spread in the timing, thereby distributing the repressive input over a wider time interval thus decreasing the sensitivity of the oscillator to perturbations. Assuming that the entrainment ability of an oscillator can be broadly characterized by its IP, our results suggest that while individual SCN neurons are all capable of generating robust sustained oscillations of the circadian TTFL, their ability to adjust to new circadian schedules varies along a ridge in the parameter space formed by the Hill coefficient and delay dispersion. Moreover, the ridge translates to a spatial structure that essentially distinguishes between the synchronized central and the more entrainable dorsal oscillator phenotypes in the SCN.

## Discussion

The time evolution of biochemical reaction networks arises from complex stochastic processes involving many species and reaction events. Inference for such systems is invariably challenged by the relative sparseness of experimental data as measurements can often only be observed for one of the participating species and at discrete time points. If collected for many cells over time the resulting data sets are of considerable size. A reduction of the full system to a model with a more modest number of parameters that can feasibly be identified from data and which retains biological interpretability is of significant importance. This can be achieved for oscillatory dynamics resulting from TTFLs by the introduction of distributed delay times which account for the natural variability in timing of unobserved processes in the larger and more complex biochemical reaction network. Inference for the resulting stochastic differential equation is subject to ongoing research. Here the filtering approach developed by [5] is further extended through Bayesian hierarchical modelling comprising a spatial component, that essentially addresses the correlation of parameters describing oscillators in nearby locations. We propose a novel measure of oscillatory robustness that allows us to estimate the posterior probability that an oscillator is inherently subject to limit cycle dynamics in contrast to noise-driven. The approach is of particular interest for modelling spatially distributed oscillators such as molecular clocks in SCN neurons. The idea that noisy systems are more easily entrained to an external input has been investigated both theoretically [35] and experimentally [21]. In particular, the latter studied the role of intrinsic noise in the SCN, and provided experimental evidence from a *Bmal*1-null mutant mice that noise and extracellular signalling are sufficient to produce oscillations when the TTFL is disrupted. Our approach extends over existing work in that it provides a framework to perform inference on the spatial distribution of the model parameters and thus to gain insight into such stochastic biochemical oscillators and their synchronisation on the basis of real experimental data. We find spatial differences across the SCN tissue, primarily in the Hill coefficient and entropy of the delay distribution and these findings are consistent across three biological replicates. In general, we identify higher values for both Hill coefficient and delay dispersion in centrally located tissues and lower values in dorsal tissues. We show how simultaneously varying the two parameters gives rise to differences in the transcriptional dynamics in cell-autonomous oscillators, represented by the IP. We thus find that the distributed delay TTFL model can separately capture both *robustness* and *entrainability* of circadian oscillations and thus has dynamics rich enough to investigate fundamental aspects of the mammalian clock, while being amenable to statistical parameter inference.

## Materials and methods

### Bioluminescense recording and preprocessing

Luciferase-reported *Cry1* expression was recorded over 5 to 6 days in three biological replicates of organotypic bilateral SCN slices using an EM-CCD camera with exposure time 0.5h resulting in ca. 250 frames of 400 by 240 pixels [3]. To achieve a signal-to-noise ratio that approximates that of the single cell level and noting that the size of a neuron corresponds to approximately 8 by 8 pixels, we aggregate the raw data into 4 by 4 pixel blocks, which we refer to as *locations*. As the proposed methodology is computationally costly we analyze a sub-sample consisting of alternate rows and columns of locations. A further de-trending is often required in the analysis of circadian bioimaging data due to consumption of luciferin substrate and is implemented in standard software for analysis of circadian data, e.g. Biodare2 (https://biodare2.ed.ac.uk/) [44]. In our analysis, at each location a (decreasing) linear trend fitted by least squares is subtracted.

### MCMC algorithm and prior distributions

This modelling framework is extended to a spatially distributed population of oscillators by placing the single-cell model in a Bayesian hierarchical structure wherein a random effects parameter model accounts for cell heterogeneity driven by extrinsic noise and phenotypic variation. Considerable statistical strength for inference can be gained generally by modelling the dependence between parameters of spatially nearby locations and we find the hierarchical model structure enables joint inference for large spatio-temporal gene expression data sets. Such data consist of indirect and noisy measurements of population sizes of chemical species through measurement processes involving, for example, bioluminescent or fluorescent reporter protein. A third layer of our hierarchical model explicitly models both measurement noise and temporal smoothing associated with such measurements. To infer parameters of the hierarchical model from spatio-temporal experimental data we design a Markov chain Monte Carlo (MCMC) algorithm which uses efficient parallelism facilitated by the imposed spatial dependency structure.

Parameter estimation is achieved through the development of a Markov chain Monte Carlo (MCMC) algorithm which enables joint inference for entire neuronal circuits.

While other distributional assumptions can be made regarding the delay distribution, the Gamma density possesses an asymmetric shape which generalises the exponential density arising exactly under restrictive modelling assumptions, and in our experience is simple and flexible enough to account for the distribution of time of intermediate unobserved processes such as translation, dimerization and nuclear export/import with spatially varying parameters.

The assumption of Gaussianity for the measurement error leads to computational advantages in the form of being able to invoke efficient filtering procedures. A residual analysis was performed as an *a posteriori* means of assessing the validity of this assumption.

The prior distribution of the multiplicative random effects is given by a Gaussian conditional autoregressive model [2]. The assumed neighbourhood structure implies a first-order Markov random field, specifically that the random effect for a given parameter at location *i* is a priori dependent on the random effects of the same parameter associated with the 8 surrounding locations. This structure is encoded by an adjacency matrix *W* with element *w*_*i,j*_ taking the value 1 if *i* and *j* are neighbouring locations and 0 otherwise. Let *w*_*i*,+_ denote the *i*^*th*^ column sum *W*, i.e. the number of neighbours associated with location *i* and let *ϵ*_*−i*_ denote the full set of random effects except *ϵ*_*i*_. The conditional prior distribution of a random effect at location *I* is given by

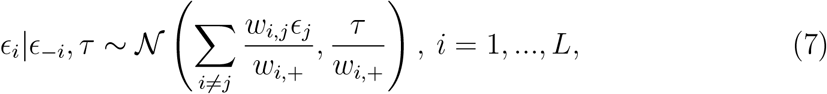

which implies an improper (non-integrable) joint prior distribution over all random effects of a given parameter. For identifiability a sum-to-zero constraint is imposed on each set of random effects. The posterior distribution resulting from components (i)-(iii) is intractable, hence we design a Markov chain Monte Carlo (MCMC) algorithm to draw a large number of samples from it. The dependence structure induced by the Markov random field essentially determines the feasibility of the computational approach to inference as the distributions involved can be evaluated in parallel over sets of non-neighbouring locations.

Prior distributions for the hyper-variances of the random effects *τ* are taken to be (moderately vague) zero-mean Gaussian with variance 25. Prior distributions for the spatial means and light scaling, *κ*, also are vague, zero-mean Gaussian with variance 100 for logarithms of *R, K* and *n* and uniform on [0, 24] and [0, 20] for the mean and standard deviation of the delay distribution respectively. The informative prior for the log degradation rate is Gaussian and informative with mean −0.55 and variance 0.25^2^, as elicited from estimates of functional half-life of luciferase [42]. We are able to obtain maximum likelihood estimates of the measurement error dispersion at each location under the assumption that it is dominated by Gaussian serially uncorrelated noise. The variance of the read noise, arising in the digitization of recorded photoelectrons, of a specific CCD camera setup can generally be estimated using a reference dark recording [15]. In the absence of such a recording it is still possible to estimate the noise variance under assumptions that two separate recordings are available, that read noise dominates other sources of noise, and that the noise process is stationary. Consider RGB video data where 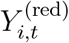 and 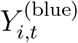 are the recorded signals in the red and blue channel at time *t* and location *i*, respectively. Under the assumption that they contain the same underlying signal and are corrupted with independent realizations of a Gaussian noise process with variance 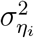 then

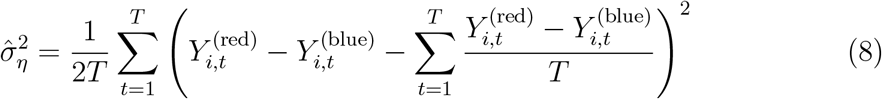

is the maximum likelihood (ML) estimator of 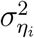. To obtain an informative prior distribution we apply this estimator to all locations and take logarithms and hence obtain an empirical distribution to which a Gaussian is fitted with mean −5.3 and standard deviation 0.17. This provides a lognormal prior distribution for the noise variance.

As the posterior distribution of the parameters is intractable we resort to designing a blocked random walk Metropolis algorithm to sample the posterior distribution. The algorithm is implemented in Matlab 2019a [26]. While computationally expensive, it is executable on a standard modern desktop PC, in large part due to two design features: the location-specific likelihoods are computed in parallel which we found to speed up computation linearly with the number of available processor cores. In addition, we make use of adaptive MCMC [30], i.e. a scheme to optimize the variances of the proposal distribution which substantially improves mixing and reduces the number of iterations required to achieve convergence. Parameter blocks are updated in a fixed scan Gibbs fashion. The blocking strategy is utilized in order to make use of the empirical covariance structure of the posterior wherever computationally feasible. The blocks are (with corresponding dimension) hyper-variances *τ* (5), global means *θ* (5), five blocks of random effects associated with each of the global means *ϵ* (L), degradation rate *µ* (1), light scaling *κ* (1) and measurement error standard deviation *σ*_*η*_ (1). For the 5-dimensional block proposals we use the estimated posterior covariances, scaled by an adaptive coefficient to obtain acceptance rates close to 0.234 and for the 1-dimensional a rate of 0.45 [29]. For each of the random effect blocks, proposals are spherical Gaussian with variances scaled using another adaptive coefficient to obtain the an acceptance rate of 0.234. While there is covariance structure in the posterior distribution of random effects that could in principle be exploited, it is prohibitively expensive in practice due to the dimension of the these parameter blocks. The adaptive coefficient *γ*^(*k*)^ for each block at iteration *k* is tuned by examining the empirical acceptance rate of the chain and taking *γ*^(*k*+1)^ = *γ*^(*k*)^ *· c*_*k*_ if the observed acceptance rate is above the target value and *γ*^(*k*+1)^ = *γ*^(*k*)^ *·* (1 *− c*_*k*_) if below. Furthermore, diminishing adaption is implemented by taking *c*_1_ = 0.02 and *c*_*k*_ = *c*_*k−*1_*−c*_*k−*1_/10^4^. The resulting chains have acceptance rates close to optimal.

As the output of an algorithm with random walk proposals typically suffers from positive autocorrelation we evaluate the output using lugsail batch means effective sample sizes [39] which calculates the equivalent sample size free of serial correlation. The algorithm is run for 6×10^4^ iterations for each experimental replicate on an 3.40 GHz Intel Core i7-6700 processor utilizing four cores for parallel evaluation of the likelihood approximation. The first half of the output is treated as burn-in and discarded and the remaining chains are used for further analysis. The computational cost is approximately 500 hours per replicate and the univariate effective sample size is *>* 100 for all parameters and locations.

### Stability criteria of macroscopic rate equation

To obtain stability criteria for the macroscopic rate equation, given by

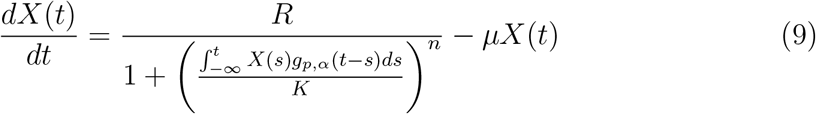

we make use of the linear chain trick [33] to obtain the following system of ODEs for *X*_0_(*t*) := *X*(*t*) and intermediate states *X*_*i*_,

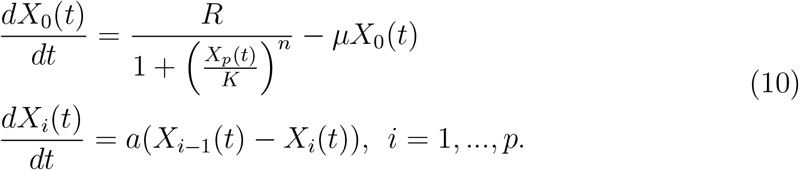

which is a monotone cyclic feedback system (MCFS) [25], i.e. flows through the system are unidirectional as *dX*_*i*_*/dt* is monotonically forced by *X*_*i−*1_. By Theorem 4.1 of ref. [25] the omega limit sets of solutions to a MCFS with negative feedback, and det(*−J* (*x*^*∗*^)) *>* 0, where *J* is the Jacobian of the system and *x*^*∗*^ is the unique fixed point *X*_*i*_ = *x*^*∗*^ s.t. *dX*_*i*_*/dt* = 0 for *i* = 0, …, *p*, are either non-constant periodic orbits or the unique fixed point. To show that det(*−J* (*x*^*∗*^)) *>* 0, we make use of the sparse structure of the (negative) Jacobian,

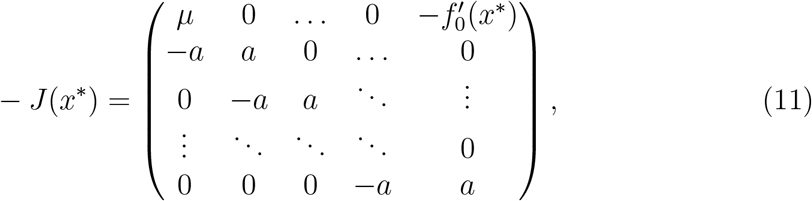

and note that det(*−J* (*x*^*∗*^)) can be decomposed as

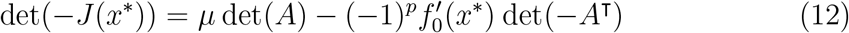

where *A* is the *p × p* bidiagonal sub-matrix obtained when deleting the first row and column of *−J* (*x*^*∗*^). The determinant of *A* is *a*^*p*^, and hence we obtain

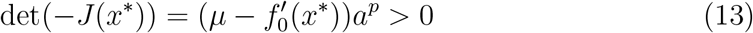

for *µ, a >* 0 and 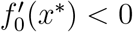 To determine whether a given set of parameters imply limit cycle or damped dynamics we examine the real part of the dominant eigenvalue of *J* (*x*^*∗*^), where a positive real part implies a limit cycle and a negative real part (and non-zero imaginary part) imply damped oscillations. To calculate the posterior probability of a limit cycle the matrix *J* is constructed for the thinned MCMC output and the real part of the largest eigenvalue is evaluated using an indicator function that takes the value 1 if the eigenvalue has positive real part and 0 if negative. The mean of the resulting chain of indicators is reported as the posterior probability of a limit cycle. Note however that the stability result is derived under *p∈* ℕ, while the MCMC samples of *p* take values in ℝ. To approximate the Jacobian we device a rounding scheme for *p* and *a* that preserves the dispersion of the delay distribution, given by

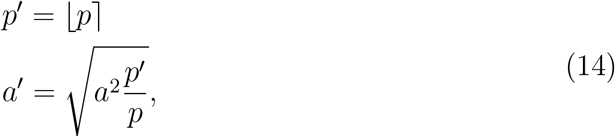

Where ⌊·⌉ denotes rounding to nearest integer. To investigate sensitivity to the rounding we repeated the analysis for ⌊*p*⌋ and ⌈*p*⌉ and conclude that the choice of rounding scheme did not impact the findings.

### Likelihood approximation of TTFL model

The likelihood of the state space model model given in Eq (1-2) is intractable, but can be approximated with the filtering procedure for distributed delay CLEs of Calderazzo et al. [5]. In essence, filtering is accomplished via a Kalman update, following a first order linearisation of the nonlinear functions involved in the mean and variance equations of the process, which are analogous to the extended Kalman-Bucy filter (EKBF) algorithm for non-delayed systems [24, 32]. Consider a time discretization of Eq (1) and (2), given by

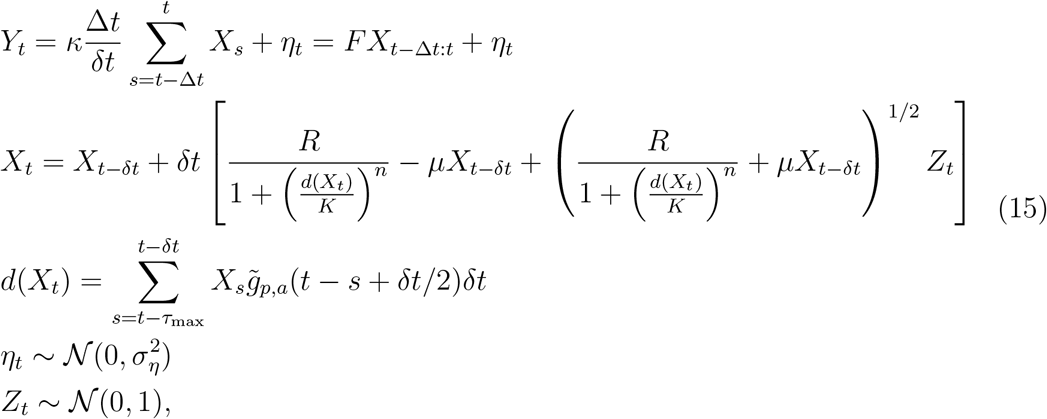

where 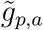 is the gamma delay density truncated at *τ*_max_. Assume there exists an optimal estimate of the normal distribution of the initial state condition given observations 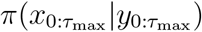 with mean vector 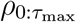 and covariance matrix 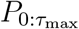 In practice we obtain these by scaling initial data by the current *κ*. By Taylor expanding the discretized state transition equation about the mean vector 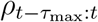 at each iteration and dropping terms of order (*δt*)^2^, we obtain the following equations for the mean and covariance

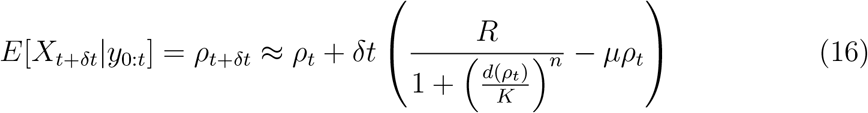

and

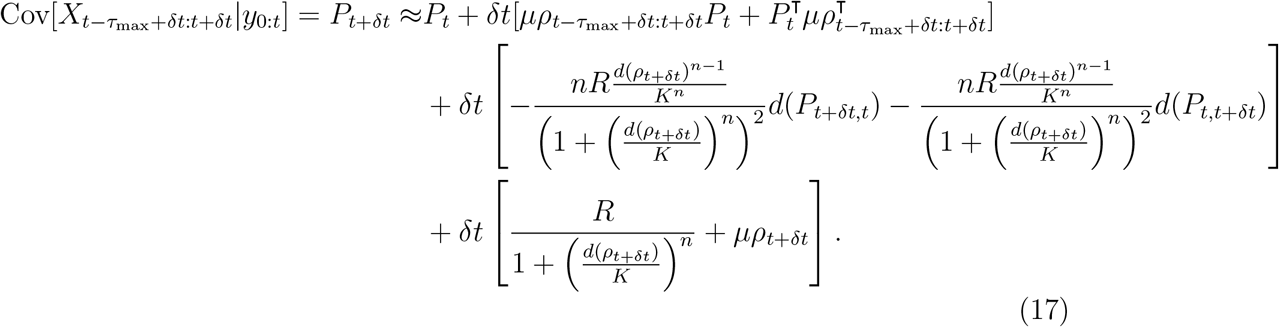

As observations are available at discrete time-points Δ*t*, 2Δ*t*, …, *T* we can use Eq (16-17) to propagate the estimates of the mean and covariance of the unobserved states up until the next observation *y*_*t*+Δ*t*_ and subsequently condition on *y*_*t*+Δ*t*_ using Kalman update

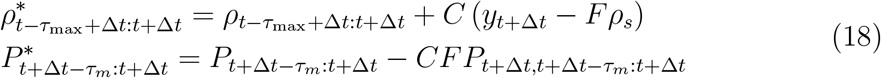

where

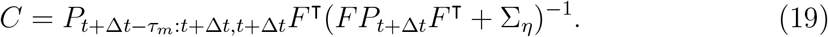

The marginal likelihood of parameters *θ* given observations can be decomposed as 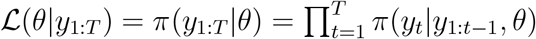 We have Gaussian errors and 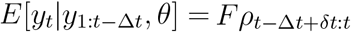 and 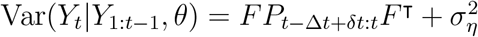. Letting 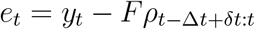 we can write the marginal log-likelihood, up to an additive constant, as

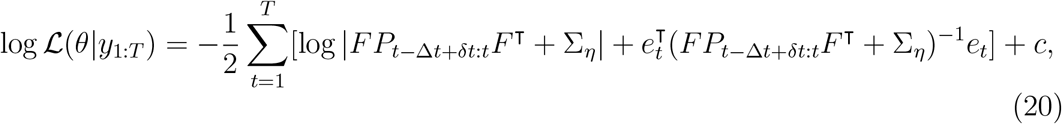

where the resulting sum is interpreted as a function of the parameters given the fixed observations.

## Ethics statement

All mouse-based work was conducted under the UK Animals (Scientific Procedures) Act 1986, with UK Home Office Licence PPL 70.8090, and local ethical approval by the LMB Animals Welfare and Ethical Review Body.

## Acknowledgements

We thank Silvia Calderazzo for valuable discussions on the methodology and its implementation. MU was supported by the ESRC (award ref. 1791198). AMJ was partially supported by the EPSRC (grants EP/R034710/1 and EP/T004134/1), and the Loyd’s Register Foundation Programme on Data-Centric Engineering at the Alan Turing Institute.

## Code availability

All Matlab code and documentation required for the analysis are available at https://github.com/mansunosson/SCN_analysis.

